# Correct laminar positioning in the neocortex influences proper dendritic and synaptic development

**DOI:** 10.1101/229948

**Authors:** Fanny Sandrine Martineau, Surajit Sahu, Vanessa Plantier, Emmanuelle Buhler, Fabienne Schaller, Lauriane Fournier, Geneviève Chazal, Hiroshi Kawasaki, Alfonso Represa, Françoise Watrin, Jean-Bernard Manent

## Abstract

The neocortex is a six-layered laminated structure with a precise anatomical and functional organization ensuring proper function. Laminar positioning of cortical neurons, as determined by termination of neuronal migration, is a key determinant of their ability to assemble into functional circuits. However, the exact contribution of laminar placement to dendrite morphogenesis and synapse formation remains unclear. Here we manipulated the laminar position of cortical neurons by knocking down Dcx, a crucial effector of migration, and show that misplaced neurons fail to properly form dendrites, spines and functional glutamatergic synapses. We further show that knocking down Dcx in properly positioned neurons induces similar but milder defects, suggesting that the laminar misplacement is the primary cause of altered neuronal development. Thus, the specific laminar environment of their fated layers is crucial for the maturation of cortical neurons, and influences their functional integration into developing cortical circuits.

## INTRODUCTION

Histogenesis of the neocortex relies on intricate developmental events eventually leading to the formation of a laminated, six-layered structure. The laminar distribution of neocortical neurons is closely linked with their ability to assemble into appropriate circuits, thereby ensuring proper cortical function. Accordingly, altered laminar positioning resulting from neuronal migration disorders is associated with clinical symptoms such as epilepsy and intellectual disability (1 – 3).

Although neocortical cytoarchitecture has long been described, developmental mechanisms underlying the establishment of neuronal connectivity remain partly unknown. Once generated from progenitors, neocortical neurons undergo extensive migration before they settle into their target layers. While migrating, neurons extend two types of processes: a leading process at their front end, and a trailing process behind. Although the trailing process is already defined as a growing axon (4 – 6), the leading process will become the apical dendrite only after migration has ended (7, 8). Therefore, a tight temporal and functional coupling exists between the eventual positioning of neocortical neurons defined by termination of migration, and the subsequent morphogenesis of dendrites. Pyramidal neurons of the neocortex are highly polarized cells, displaying apical and basal dendritic arbors. Synapses being formed at precise locations on dendritic spines, the size and shape of dendritic arbors are key determinants for the number and distribution of synaptic contacts. However, the exact contribution of laminar positioning to dendrite morphogenesis and synapse formation remains unclear.

In the present study, we investigated whether and how a laminar misplacement may influence the morphological and functional maturation of neocortical neurons. We induced altered laminar positioning through in utero knockdown of doublecortin (Dcx), a well-known effector of migration, and analyzed how ectopic neurons developed postnatally. We observed significant changes in dendrite and spine morphologies accompanied by altered glutamatergic synaptic transmission. We also evaluated the relative contributions of laminar misplacement and Dcx to these changes. Our data indicate that, besides a mild influence of Dcx on neuronal maturation, ectopic neuronal positioning is a major contributor to the altered dendritic morphogenesis and impaired development of functional glutamatergic synapses.

## RESULTS

### Ectopic neurons fail to develop proper dendritic arbors and spines

To study whether a correct laminar positioning may influence the morphological and functional maturation of neocortical neurons, we disturbed neuronal migration by knocking down Dcx *in utero*. At embryonic day 15 (E15), a plasmid encoding a short hairpin RNA (shRNA) targeting Dcx was electroporated in the cortical progenitors of rat embryos (9) (Figure 1A). Co-electroporation of a plasmid encoding the red fluorescent protein mCherry allowed visualization of the electroporated neurons in post-natal stages. At post-natal days 14-15 (P14-15), mCherry+ neurons electroporated with Dcx shRNA had failed to migrate and were located ectopically within the white matter, as previously described. By contrast, in control brains electroporated with a Dcx-mismatch shRNA, mCherry+ neurons migrated properly into layer V (Figure 1B).

**Figure 1:**
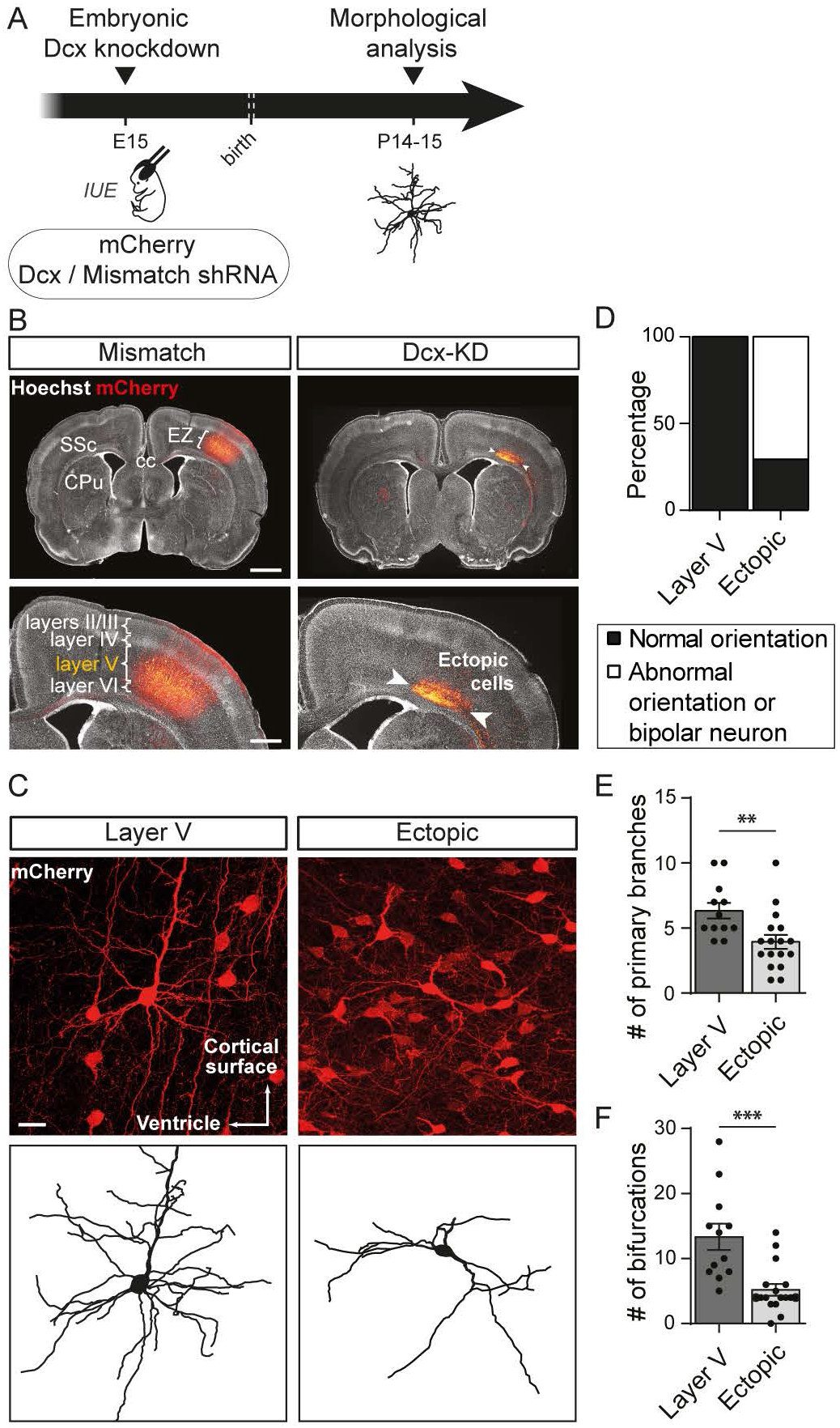
Altered neuronal positioning impairs neuronal polarity and morphology. (A) Schematic of the experimental timeline. Embryos were electroporated *in utero* at E15 with a plasmid encoding a short hairpin RNA (shRNA) targeting Dcx 3’UTR to create an embryonic Dcx knockdown or with a mismatch shRNA for control. A second plasmid encoding a red fluorescent protein, mCherry, was co-electroporated to spot the electroporated neurons. *IUE: in utero* electroporation (B) Representative neocortical sections showing that mismatch neurons are properly placed in layer V of the somato-sensory cortex (SSc) while Dcx-KD neurons ectopically between the corpus callosum (cc) and layer VI of the cortex at P14. EZ = electroporation zone; CPu = caudate putamen. Top panels, scale bar = 2 mm. Bottom panels, scale bar = 1 mm (C) High magnification confocal images of layer V and ectopic neurons used to reconstruct their dendritic arbors. The 3D reconstructions are depicted below. Scale bar = 100 μm. (D) Stacked bar graph showing the percentage of normally oriented neurons and bipolar or abnormally oriented neurons in each set of data analyzed in (E) and (F) (E,F) Quantification of the mean number of primary branches (E) and bifurcations (F) in 3D reconstructions from layer V and ectopic neurons. Each dot represents a neuron. (E) t-test, p-value < 0.0074; (F) Mann-Whitney test, p-value < 0.0001

To analyze whether altered neuronal positioning has an impact on neuronal polarity and overall morphology, we imaged ectopic neurons from Dcx knockdown (Dcx-KD) brains compared to layer V neurons from mismatch brains and scored their apical dendrite orientation (Figure 1C-D). Apical dendrites of layer V neurons were 100% (12/12 neurons) radially oriented towards the cortical surface whereas less than 30% (5/17 neurons) of ectopic neurons were oriented properly. The remaining ectopic neurons were either oriented aberrantly or presented an abnormal bipolar morphology, preventing us from clearly defining an apical dendrite (Figure 1D). We then checked whether altered cortical positioning influenced dendritic growth by reconstructing the dendritic arbor and analyzing the complexity of basal dendrites. Ectopic neurons grew significantly fewer primary branches than layer V neurons (6.3 ± 0.6 vs 3.9 ± 0.5; t-test, p = 0.0074; n = 12 layer V neurons, 17 ectopic neurons) and their dendritic arbor was less ramified (13.3 ± 2 vs 5.2 ± 0.9; Mann-Whitney test, p < 0.0001; n = 12 layer V neurons, 17 ectopic neurons) (Figure 1E-F).

To evaluate the impact of altered positioning and dendritic growth on spine formation, we manually reconstructed the spines from the mCherry signal amplified by immunohistochemistry (Figure 2A). In ectopic neurons, the linear density of spines was decreased by 40% compared to control neurons in layer V (35.6 ± 3.2 vs 21.6 ± 2.6 spines / 100 μm; t-test, p = 0.0024; n = 13 neurons / condition) (Figure 2B). In addition, successfully formed spines had an altered morphology including wider heads (0.498 ± 0.005 vs 0.535 ± 0.01 μm; Mann-Whitney test, p < 0.0001; n = 1953 spines from 13 layer V neurons and 501 spines from 13 ectopic neurons) and wider necks (0.27 ± 0.002 vs 0.39 ± 0.009 μm; Mann-Whitney test, p < 0.0001; n = 1953 spines from 13 layer V neurons and 501 spines from 13 ectopic neurons) although their length was comparable to spines from layer V neurons (1.281 ± 0.015 vs 1.281 ± 0.034 μm; Mann-Whitney test, p = 0.15; n = 1953 spines from 13 layer V neurons and 501 spines from 13 ectopic neurons) (Figure 2C-E). Together, these data suggest that an appropriate positioning ensures proper cell polarity, dendritic growth and spine morphology.

**Figure 2:**
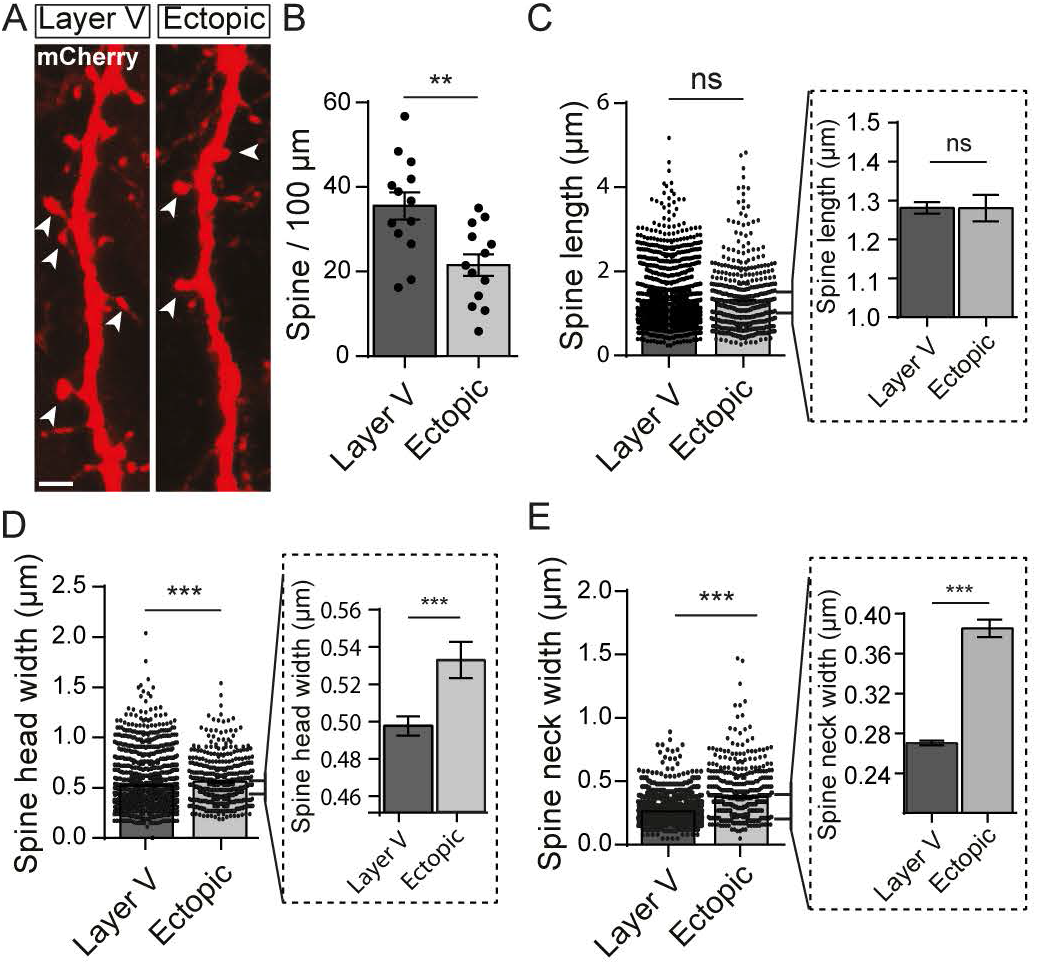
Ectopic neurons display a reduced spine density and altered spine morphology. (A) High magnification confocal images of dendrites from layer V and ectopic neurons electroporated with mCherry. Representative dendritic spines are indicated by arrow heads. Scale bar = 2 μm (B) Bar graph illustrating the mean number of spines per 100 μm of dendrite. Each dot represents a neuron. T-test, p-value = 0.0024 (C-E) Quantification of the mean dendritic spine length (C), head width (D) and neck width (E) in layer V and ectopic neurons. Each dot represents a spine. Bar graphs in dotted line frames are higher magnifications of the initial bar graphs. (D, E) Mann-Whitney test, p-value < 0.0001

### Glutamatergic synaptogenesis is altered in ectopic neurons

Because dendritic spines are the sites of most excitatory synapses, the spine alterations reported above could be associated with impaired glutamatergic synaptogenesis. To check this point, glutamatergic post-synapses were genetically labelled with PSD-95-GFP, a scaffolding protein that anchors glutamate receptors and had been fused with GFP. We co-electroporated a floxed-stop-PSD-95-GFP plasmid at E15 with plasmids encoding Dcx shRNA, mCherry and a tamoxifen-inducible Cre-recombinase. The Cre-recombinase was activated at P1 by intraperitoneal injection of tamoxifen to induce PSD-95-GFP expression (Figure 3A). A single tamoxifen injection induced a low level of recombination and allowed for a sparse labelling of PSD-95-GFP, making it easier to follow dendrites of individual neurons (Figure 3B). Once expressed, PSD-95-GFP proteins formed puncta localized mainly on spine heads, where the post-synapse is usually assembled (Figure 3C). Electron microscopy experiments confirmed that PSD-95-GFP was localized in spines near the electron dense post-synaptic density (PSD) facing pre-synaptic elements containing vesicles (Figure 3D). We then quantified the linear density of PSD-95-GFP^+^ puncta along dendritic segments localized within 5-80 μm of the soma. In ectopic neurons, the PSD-95-GFP^+^ mean density was significantly lower than in layer V neurons (87.54 ± 4.03 vs 15.31 ± 2.53 puncta / 100 μm; t-test, p < 0.0001; n = 13 neurons / condition) (Figure 3E-F), suggesting that ectopic neurons form less glutamatergic synapses. In addition, spines containing PSD-95 are generally classified as more differentiated (73) so we analyzed the proportion of those containing PSD-95-GFP^+^ puncta in ectopic and layer V neurons. In ectopic neurons, about 90% (180/217) of spines analyzed were devoid of PSD-95-GFP^+^ puncta (n = 6 neurons) whereas this fraction was only 24% (199/621) in layer V neurons (n = 6 neurons) (Figures 3E and 3G). Although we did not quantify it, we noted that PSD-95-GFP^+^ puncta were mostly localized on the shaft of dendrites in ectopic neurons. Thus, in addition to a decreased spine density, ectopic neurons form less spines with a mature PSD scaffold.

**Figure 3:**
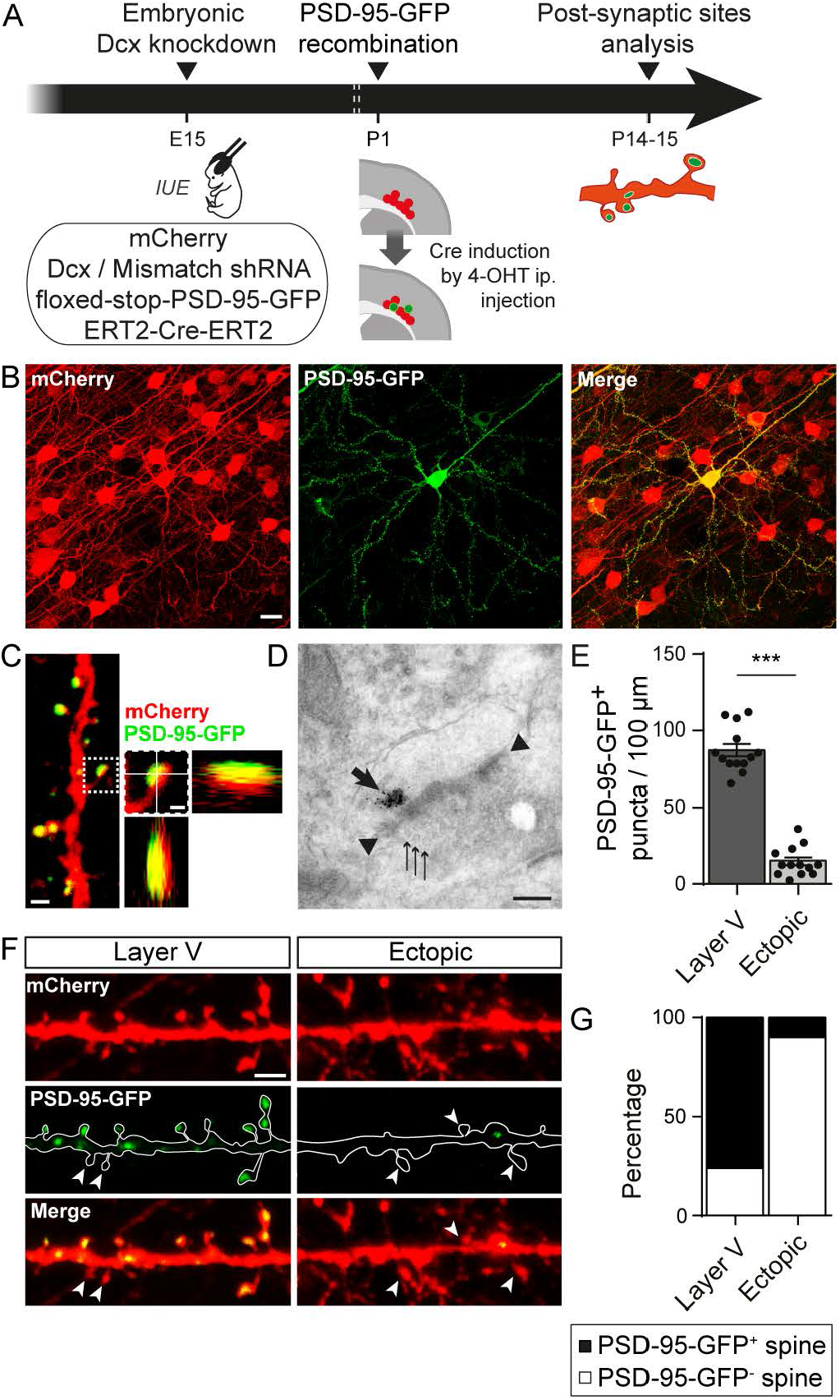
Altered neuronal positioning decreases the density of glutamatergic synapses. (A) Animals were electroporated at E15 with three plasmids encoding for the fluorescent marker mCherry, a Dcx shRNA and a tamoxifen activable Cre-recombinase (ERT^2^-Cre-ERT^2^). A fourth conditional plasmid was co-electroporated to express the post-synaptic glutamatergic protein PSD-95-GFP (floxed-stop-PSD-95-GFP). Sparse recombination of the floxed-stop-PSD-95-GFP plasmid was induced in a subset of electroporated neurons through a single intraperitoneal (ip.) injection of 4-hydroxytamoxifen (4-OHT) at P1. Glutamatergic post-synaptic sites were analyzed at P14-15. *IUE: in utero* electroporation (B) Confocal images of the electroporated zone (red) with a single neuron having undergone recombination of the floxed-stop-PSD-95-GFP plasmid (green) following 4-OHT injection. Scale bar = 20 μm (C) High magnification confocal image of a dendrite with PSD-95-GFP^+^ spines. On the right, higher magnification of one dendritic spine and its orthogonal views show the PSD-95-GFP^+^ punctum colocalize with the mCherry filled spine. Left image scale bar = 1 μm; right image scale bar = 0,5 μm (D) Electron microscopy image of a PSD-95-GFP^+^ post-synapse facing a presynaptic compartment containing vesicules. The thick arrow shows immunogold labelled PSD-95-GFP proteins; arrow heads show electron-dense thickening of the post-synaptic membrane; thin arrows show pre-synaptic vesicules. Scale bar = 0,2 μm (E) Bar graphs and scatter dot plots illustrating the density of PSD-95-GFP^+^ puncta per 100 μm of dendrites in layer V and ectopic neurons. t-test, p-value < 0.0001 (F) Confocal images of mCherry+ dendrites and PSD-95-GFP^+^ puncta from layer V and ectopic neurons. Arrow heads indicate spines devoid of PSD-95-GFP^+^ puncta. Scale bar = 2 μm (G) Stacked bar graphs showing the relative percentages of PSD-95-GFP^+^ and PSD-95-GFP^−^ spines in layer V and ectopic neurons.

To investigate whether a reduced density of spines and a reduced PSD scaffold translate into altered synaptic transmission, we recorded miniature excitatory currents (mEPSC) from electroporated neurons (Figure 4A-B). We successfully recorded mEPSCs in ectopic neurons, indicating that these cells received synaptic inputs. However, cumulative frequency curves of mEPSC inter-event intervals revealed a significant shift towards higher values, suggesting a decreased frequency of mEPSCs in ectopic neurons as compared to layer V neurons (Kolmogorov-Smirnov test, p < 0.0001; n = 8114 events from 15 layer V neurons, 5364 events from 18 ectopic neurons) (Figure 4C). This decreased frequency of mEPSCs is in accordance with our previous results suggesting that ectopic neurons form less synapses. Additionally, we analyzed the amplitude and half-width of events occurring in ectopic neurons and compared them to those of layer V neurons. The cumulative frequency curve of mEPSC amplitude was shifted towards higher values in ectopic neurons, indicating an increased amplitude of mEPSCs in this population (Kolmogorov-Smirnov test, p < 0.0001; n = 1348 events from 14 layer V neurons, 1221 events from 18 ectopic neurons) (Figure 4D). The half-width curve showed a minor, although significant, shift towards smaller values (Kolmogorov-Smirnov test, p < 0.0001; n = 1348 events from 14 layer V neurons, 1221 events from 18 ectopic neurons) (Figure 4E). Overall, our data suggest that migration into the appropriate layer is important for dendritic growth, spine formation and glutamatergic synaptogenesis, both on a morphological and functional level.

**Figure 4:**
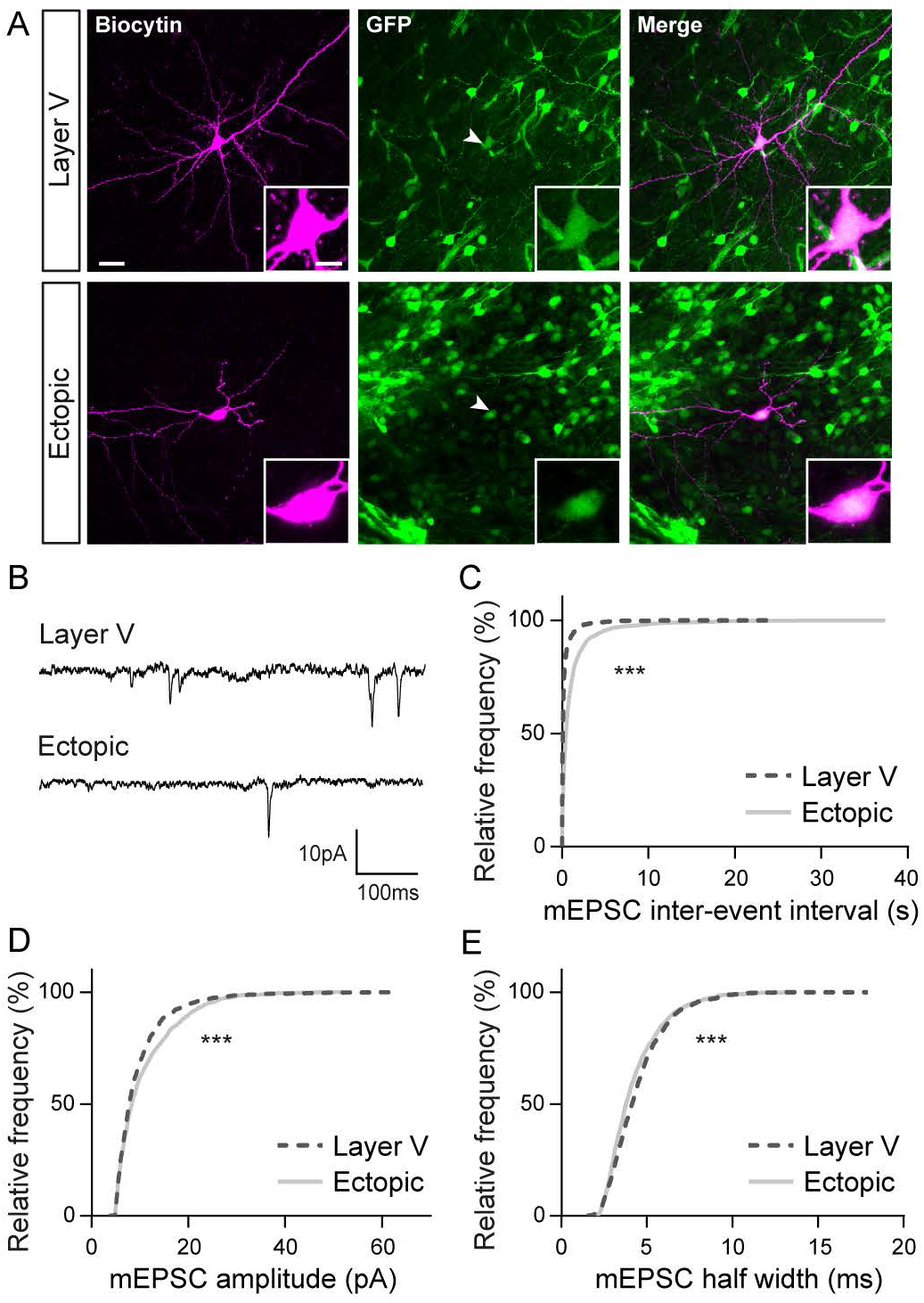
Miniature glutamatergic currents are impaired in ectopic neurons. (A) Confocal images of layer V and ectopic neurons filled with biocytin during electrophysiological recordings. Recorded cells were electroporated neurons as indicated by their GFP^+^ fluorescence (arrow heads). Inserts show higher magnification images of the somas. Scale bars = 30 μm / 10 μm (B) Representative traces of the miniature excitatory post-synaptic currents (mEPSCs) recorded from layer V and ectopic neurons and analyzed in (C-E) (C-E) Cumulative distributions of the inter-event interval (C), amplitude (D) and half width (E) values of mEPSCs from layer V and ectopic neurons. Kolmogorov-Smirnov test, p-value < 0.0001

### Dcx impacts spine morphology and synaptogenesis

The experimental paradigm we used relies on Dcx knockdown to induce ectopic positioning of neurons. Therefore, we cannot rule out that the developmental defects we observed were directly due to Dcx knockdown, independently of the position of neurons. For this reason, it was important to check if Dcx plays a role in dendritic growth, spine formation or glutamatergic synaptogenesis.

Bai *et al*. previously reported(9) that non-electroporated neurons are present amongst the ectopic cells aggregated within the white matter (Figure 5A-B). Although these cells are ectopic, they expressed Dcx during their development (“ectopic Dcx^+^ neurons”) unlike electroporated neurons (“ectopic Dcx-KD neurons”). We recorded mEPSCs from both populations to check if the development and function of glutamatergic synapses were affected by Dcx-KD (Figure 5B-C). The inter-event interval distribution was shifted towards lower values in ectopic Dcx^+^ neurons indicating a higher frequency of mEPSCs in neurons that expressed Dcx during development (Kolmogorov-Smirnov test, p < 0.0001; n = 4771 events from 10 ectopic Dcx^+^ neurons, 5364 events from 18 ectopic Dcx-KD neurons) (Figure 5D). Moreover, the cumulative frequency curve of mEPSC amplitude was slightly, but significantly, changed (Kolmogorov-Smirnov test, p = 0.0067; n = 1049 events from 10 ectopic Dcx^+^ neurons, 1221 events from 18 ectopic Dcx-KD neurons) and the cumulative frequency curve of mEPSC half-width was shifted towards higher values in ectopic Dcx^+^ neurons (Kolmogorov-Smirnov test, p < 0.0001; n = 1049 events from 10 ectopic Dcx^+^ neurons, 1221 events from 18 ectopic Dcx-KD neurons) (Figure 5E-F). Following the electrophysiological recordings, neurons were filled with biocytin and reconstructed to analyze their overall morphology (Fig 5B,G). As reported above, most ectopic Dcx-KD neurons were aberrantly oriented or presented an abnormal morphology preventing us to define an apical dendrite (10/11 neurons). In most cases, ectopic Dcx^+^ neurons were able to form an apical dendrite but were not oriented properly towards the cortical surface (6/7 neurons) (Figure 5H). Analysis of the basal dendritic arbor revealed no significant difference in the number of primary branches formed by ectopic Dcx^+^ and Dcx-KD neurons (4.29 ± 0.78 vs 2.73 ± 1.42; Mann-Whitney test, p = 0,145; n = 7 ectopic Dcx^+^ neurons, 11 ectopic Dcx-KD neurons) (Figure 5I). However, ectopic Dcx^+^ neurons were more ramified than ectopic Dcx-KD neurons, indicating that Dcx might play a role in the formation of the dendritic arbor (10 ± 2.28 vs 4.09 ± 0.73; Mann-Whitney test, p = 0,027; n = 7 ectopic Dcx^+^ neurons, 11 ectopic Dcx-KD neurons) (Figure 5J). These electrophysiological and morphological differences between ectopic Dcx^+^ and Dcx-KD neurons suggest that Dcx is involved in dendritic growth and synaptogenesis in addition to its canonic role in migration.

**Figure 5:**
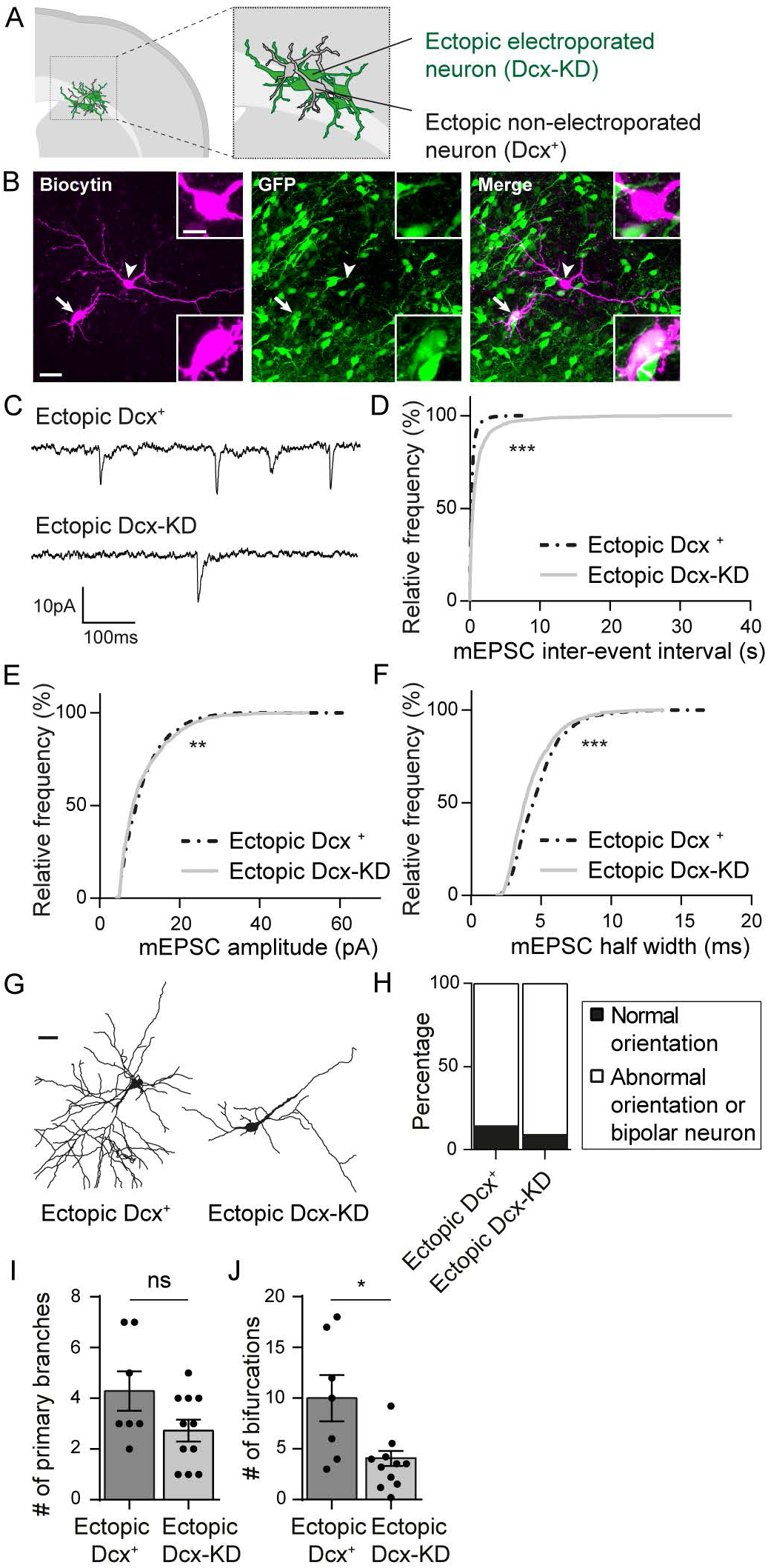
Dcx expression is important for glutamatergic synaptogenesis and dendritic growth. (A) Schematic depicting the presence of non-electroporated ectopic neurons amongst electroporated ectopic neurons. (B) Confocal images of ectopic neurons filled with biocytin during electrophysiological recordings. The arrow indicates an electroporated GFP^+^ neuron (Dcx-KD) while the arrow head shows a nonelectroporated GFP^−^ neuron (Dcx^+^). Side bar images show higher magnification images of the somas. Scale bars = 30 μm / 10 μm (C) Representative traces of mEPSCs recorded from ectopic Dcx^+^ and Dcx-KD neurons and analyzed in (D-F) (D-F) Cumulative distributions of the inter-event interval (D), amplitude (E) and half width (F) values of mEPSCs from ectopic Dcx^+^ and Dcx-KD neurons. Kolmogorov-Smirnov test, (D, F) p-value < 0.0001, (E) p-value = 0.0067 (G) 3D reconstructions of ectopic Dcx^+^ and Dcx-KD neurons used to analyze their denritic arbor in (I) and (J). Scale bar = 30 μm (H) Stacked bar graph showing the relative percentages of normally oriented neurons and abnormally oriented or bipolar neurons analyzed in (I) and (J) (I, J) Bar graphs and scatter dot plots showing the mean number of primary branches (I) and bifurcations (J) in ectopic Dcx^+^ and Dcx-KD neurons. Each dot represents a neuron. (J) Mann-Whitney test, p-value = 0.027

To further test this hypothesis, we designed a conditional knockdown strategy to silence Dcx expression after migration has ended. We electroporated at E15 a tamoxifen and Cre-dependent plasmid encoding a Dcx shRNA and induced its expression with a tamoxifen injection at P1 (Figure 6A). This post-natal knockdown (pKD) took effect after migration had ended so Dcx-pKD neurons migrated to layer V similarly to control neurons electroporated with a mismatch shRNA (Figure 6B). On the other hand, Dcx-pKD was induced early enough to study its effect on dendrite formation and early synaptogenesis, both events mostly occurring during the first post-natal weeks.

**Figure 6:**
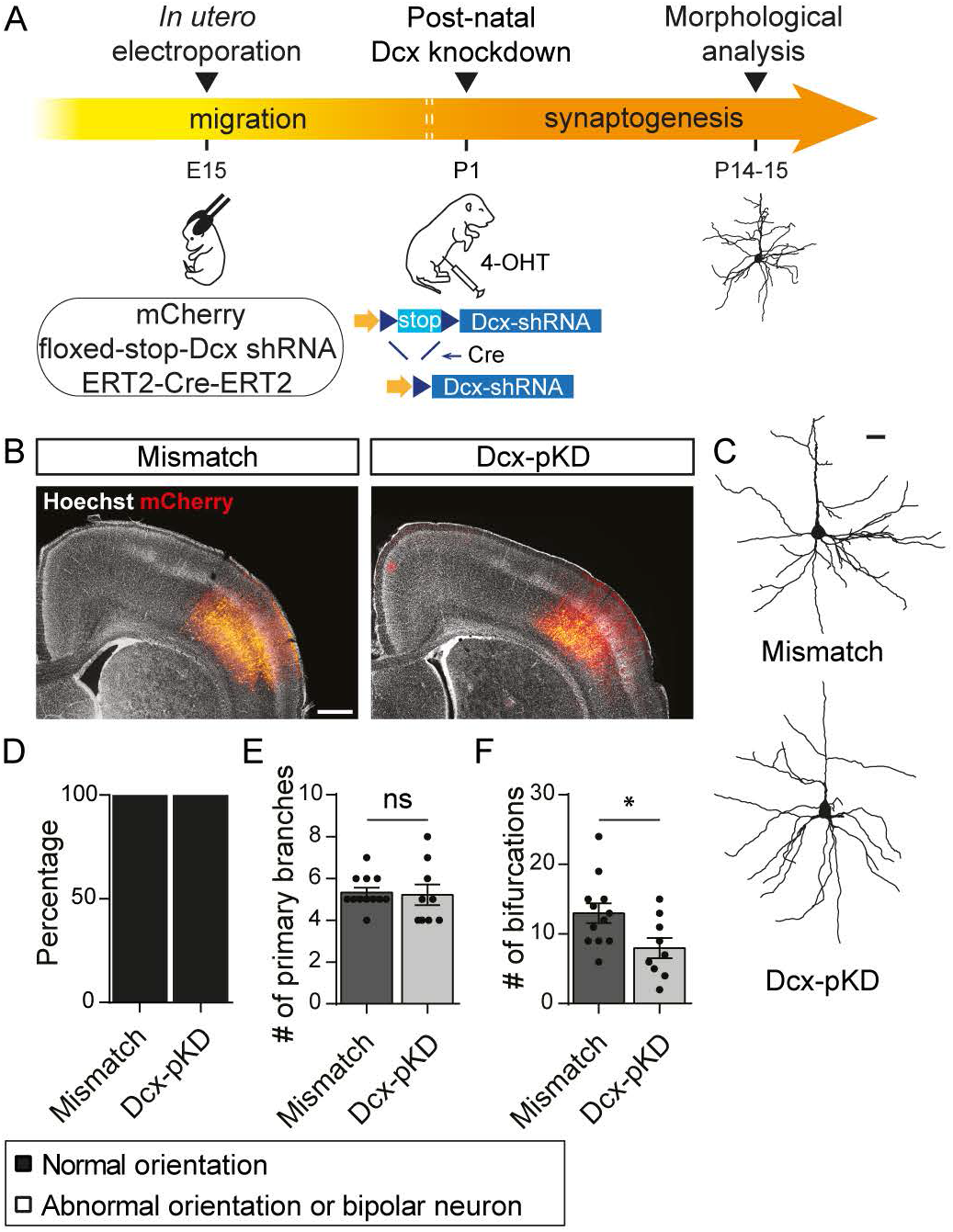
Post-natal knockdown of Dcx induces a simplication of the dendritic arbor. (A) Schematic of the experimental timeline. Rats were electroporated at E15 with a conditional plasmid encoding for a Dcx shRNA (floxed-stop-Dcx-shRNA) and two constitutive plasmids encoding mCherry and a tamoxifen-activable Cre (ERT2-Cre-ERT2). The post-natal Dcx knockdown was induced at P1 through Cre-mediated recombination following 4-OHT intraperitoneal injection. Neurons were reconstructed at P14-15 to analyze the morphology of their basal dendritic arbor. (B) Low magnification images of electroporated cortices where electroporated neurons (red) have migrated properly to layer V both in mismatch and Dcx-pKD brains. Scale bar = 1 mm (C) 3D reconstructions of mismatch and Dcx-pKD neurons used to analyze their denritic arbor in (E) and (F). Scale bar = 100 μm (D) Stacked bar graph showing the relative percentages of normally oriented neurons and abnormally oriented or bipolar neurons analyzed in (E) and (F) (E,F) Bar graphs and scatter dot plots showing the mean number of primary branches (E) and bifurcations (F) in mismatch and Dcx-pKD neurons. Each dot represents a neuron. (F) t-test, p-value = 0.026

We first analyzed Dcx-pKD effect on neuronal polarity and morphology. We observed that 100% (9/9 neurons) of Dcx-pKD neurons presented a pyramid shaped soma and their apical dendrites were radially oriented towards the cortical surface, similarly to mismatch neurons (12/12 neurons) (Figure 6C-D). Three-dimensional reconstructions of basal dendritic arbor revealed that they extended the same number of primary branches as mismatch neurons (5.3 ± 0.22 vs 5.2 ± 0.49; t-test, p = 0.826; n = 12 mismatch neurons, 9 Dcx-pKD neurons) but these branches were less ramified (13 ± 1.42 vs 8 ± 1.44; t-test p = 0.026; n = 12 mismatch neurons, 9 Dcx-pKD neurons) (Figure 6C and 6E-F). These data suggest that Dcx influences dendritic growth in properly positioned neurons, although to a lesser extent than in ectopic positions.

We then checked for a potential effect of Dcx-pKD on spine density and morphology. In Dcx-pKD neurons properly placed in layer V, spine density was similar to the one in mismatch neurons (66.8 ± 5.9 vs 56.2 ± 8.0 spines / 100 μm; t-test, p = 0.296; n = 15 neurons / condition) (Figure 7A-B). Closer analysis showed that spines from both groups also had the same length (1.308 ± 0.012 vs 1.306 ± 0.014 μm; Mann-Whitney test, p = 0.717; n = 3235 spines from 15 mismatch neurons, 2124 spines from 15 Dcx-pKD neurons). However, spines had a slightly wider head (0.468 ± 0.004 vs 0.485 ± 0.004 μm; Mann-Whitney test, p < 0.001; n = 3235 spines from 15 mismatch neurons, 2124 spines from 15 Dcx-pKD neurons) and a slightly wider neck (0.264 ± 0.002 vs 0.284 ± 0.002 μm; Mann-Whitney test, p < 0.001; n = 3235 spines from 15 mismatch neurons, 2124 spines from 15 Dcx-pKD neurons) in Dcx-pKD neurons (Figure 7C-E), consistent with a role for Dcx in spine formation.

**Figure 7:**
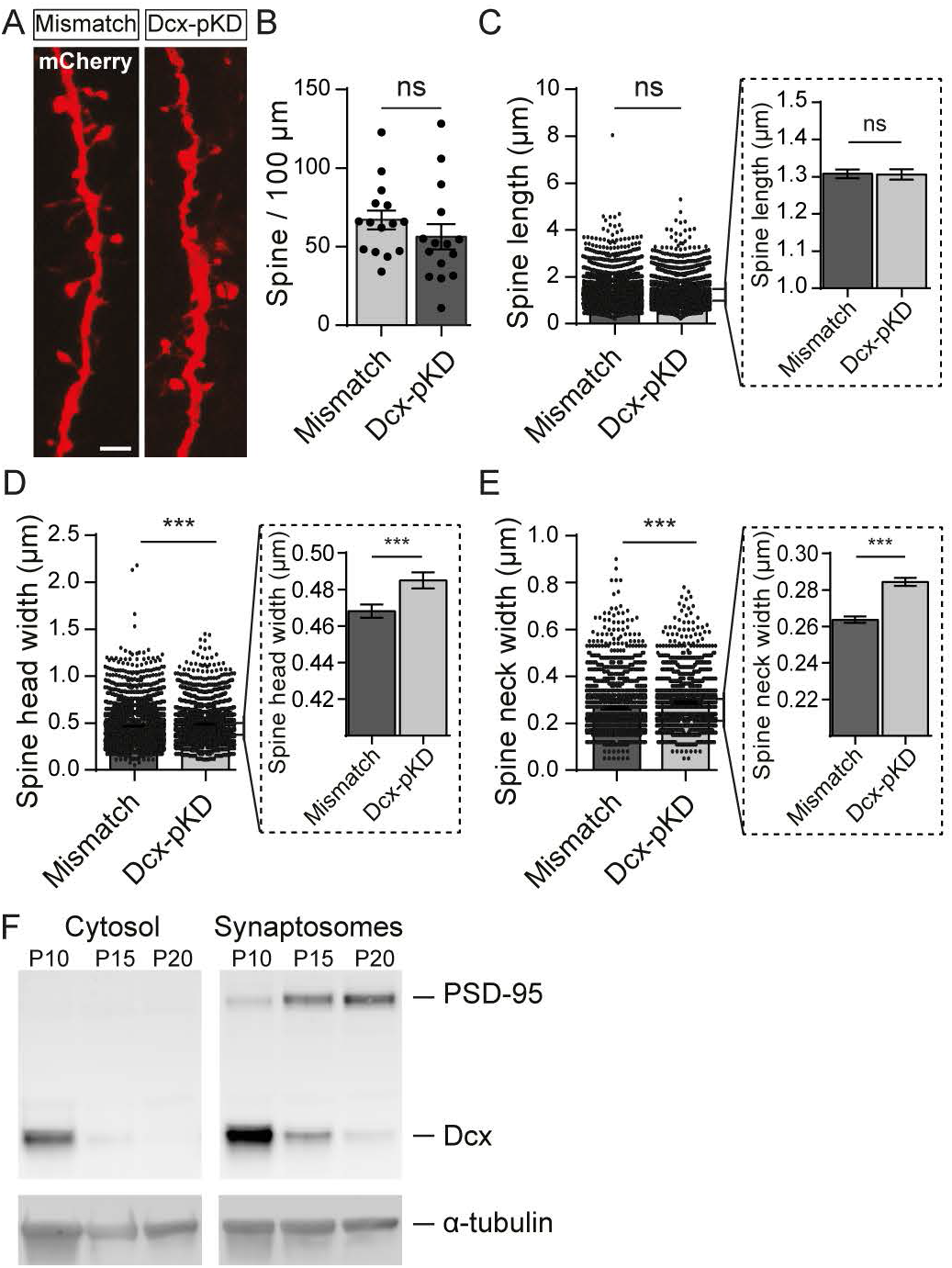
Post-natal knockdown of Dcx slightly increases dendritic spine head and neck widths. (A) High magnification confocal images of dendritic spines from mismatch and Dcx-pKD neurons coelectroporated with mCherry. Scale bar = 2 μm (B) Quantification of the density of spines for 100 μm of dendrites in mismatch and Dcx-pKD neurons. Each dot represents a neuron. (C-E) Bar graphs and scatter dot plots illustrating the mean dendritic spine length (C), head width (D) and neck width (E) in mismatch and Dcx-pKD neurons. Each dot represents a spine. Dotted line frames hold higher magnifications of the initial bar graphs. (D, E) Mann-Whitney test; p-value < 0.001 (F) Western blot of the synaptosomal and cytosolic fractions isolated from cortices of P10, P15 and P20 non-electroporated rats. (Top panels) Detection of PSD-95 (top) and Dcx (bottom). (Bottom panels) Detection of α-tubulin for control

To be consistent with this notion, Dcx should be present in the synaptic compartment during the first postnatal weeks. However, in cortical L5 neurons, very little dendritic spines are visible during this period making difficult any immunohistochemical evaluation. We therefore decided to analyze by Western blot cytosolic and synaptosomal fractions isolated from P10, P15 and P20 rat neocortex (Figure 7F) and found that Dcx was enriched in the synaptosomal as compared to the cytosolic fraction. Dcx expression was also down regulated from P10 to P20, as opposed to the PSD95 which expression increased during the same developmental period. These results agree with previous reports investigating the maturation of mouse synaptosomal proteome (10, 11).

Because Dcx seems involved in spine formation, glutamatergic synaptogenesis could also be affected by Dcx post-natal knockdown. We quantified the density of PSD-95-GFP^+^ puncta as described above and found that the linear density of PSD-95-GFP^+^ puncta was similar in both groups (101.4 ± 8.8 vs 89.1 ± 7.4 puncta / 100 μm; t-test, p = 0.296; n = 16 mismatch neurons, 15 Dcx-pKD neurons) (Figure 8A-B). Moreover, 20% (500/2913) of spines from mismatch neurons (n = 14 neurons) and 25% (490/2046) of spines from Dcx-pKD neurons (n = 15 neurons) did not contain PSD-95-GFP^+^ puncta (Figure 8C). Overall, our data show that the amount of PSD scaffold is mostly unaffected in Dcx-pKD neurons. Nevertheless, the spine alterations reported above could be associated with altered synaptic transmission. We checked this point by recording mEPSCs from Dcx-pKD neurons and mismatch neurons (Figure 8D-E). In Dcx-pKD neurons, the cumulative frequency curve of mEPSC inter-event interval was shifted towards higher values, revealing a decreased frequency of mEPSCs in Dcx-pKD neurons compared to mismatch neurons (Kolmogorov-Smirnov test, p < 0.0001; n = 8438 events from 13 mismatch neurons, 5756 events from 18 Dcx-pKD neurons) (Figure 8F). In addition, although the cumulative frequency curve of mEPSC amplitude revealed no change in Dcx-pKD neurons compared to mismatch neurons, (Kolmogorov-Smirnov test, p = 0.471; n = 1205 events from 13 mismatch neurons, 1118 events from 13 Dcx-pKD neurons), the cumulative frequency curve of mEPSC half-width was slightly but significantly shifted towards lower values in Dcx-pKD neurons (Kolmogorov-Smirnov test, p < 0.0001; n = 1205 events from 13 mismatch neurons, 1118 events from 13 Dcx-pKD neurons) (Figure 8G-H). Altogether, these data are consistent with the notion that Dcx, in addition to its known role on migration, influences spine formation, as well as the proper development of functional glutamatergic synapses.

**Figure 8:**
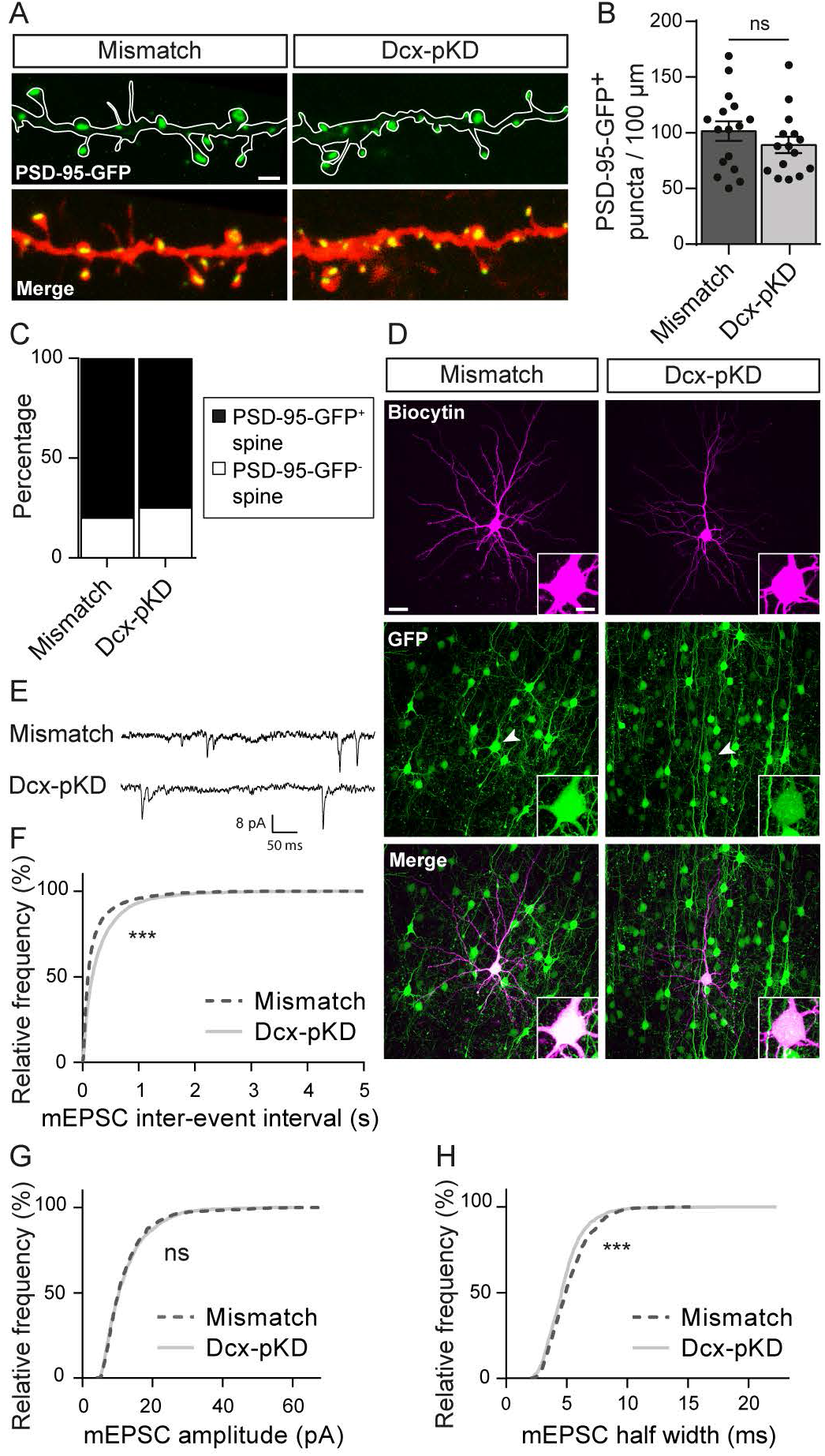
Miniature excitatory currents are impaired after post-natal knockdown of Dcx. (A) High magnification confocal images of mCherry+ dendrites and PSD-95-GFP^+^ puncta from mismatch and Dcx-pKD neurons. Scale bar = 2 μm (B) Bar graph and scatter dot plots illustrating the density of PSD-95-GFP^+^ puncta per 100 μm of dendrites in mismatch and Dcx-pKD neurons. (C) Stacked bar graphs showing the relative percentages of PSD-95-GFP^+^ and PSD-95-GFP^−^ spines in mismatch and Dcx-pKD neurons. (D) Confocal images of pyramidal neurons filled with biocytin during electrophysiological recordings. Recorded cells were electroporated neurons as indicated by their GFP^+^ fluorescence (arrow heads). Inserts show higher magnification images of the somas. Scale bar = 30 μm / 10 μm (E) Representative traces of mEPSCs recorded from mismatch and Dcx-pKD neurons and analyzed in (F-H) (F-H) Cumulative distributions of the inter-event interval (F), amplitude (G) and half width (H) values of mEPSCs from mismatch and Dcx-pKD neurons. Kolmogorov-Smirnov test, p-value < 0.0001

## DISCUSSION

In this study, we investigated whether and how laminar misplacement influences the morphological and functional maturation of neocortical neurons. We showed that ectopic positioning impairs the orientation, dendritic growth, spine development and formation of functional glutamatergic synapses in developing neurons. We also highlighted a new role for Dcx in regulating the formation of dendrites and fine-tuning glutamatergic transmission in the cortex. Accordingly, in our model of migration defect induced by Dcx knockdown, the laminar misplacement and Dcx-KD acted synergistically to alter neuronal development. However, since a post-natal knockdown of Dcx induced milder defects, we identified the laminar misplacement as the major contributor to the observed altered neuronal development.

Several rodent models and experimental strategies have been utilized to alter the laminar placement of cortical neurons and study how their maturation is impacted at unusual positions. Most studies, however, were focused on axonal projections and dendritic morphology, and rarely on the development of synapses and spines. Prenatal exposure to ionizing radiations leads to the formation of neuronal masses composed of ectopic neurons born concomitantly to irradiation. Injection of retrograde tracers revealed that these ectopic neurons develop long-distance projections to the spinal cord, as do layer V neurons born at similar gestational ages (12, 13). In genetic models with subcortical band heterotopia similar observations were made, with ectopic neurons developing normal corticospinal and corticothalamic projections (14, 15). Heterotopic transplants of embryonic neocortex combined with tracing studies confirmed these findings. Projections sent by neurons grafted at ectopic positions were similar to those of sister neurons sharing the same site of origin (16). Together, these observations suggest that migration of cortical neurons to their fated layers may not be essential for growing axons to reach appropriate targets. On the contrary, neuronal polarity and dendritic outgrowth were both severely affected in the same rodent models. Ectopic neurons in brains prenatally exposed to irradiation or chemicals displayed abnormally oriented apical dendrites (13, 17 – 19). Randomly oriented apical dendrites were also described in neurons grafted at ectopic locations (20). Similarly misoriented neurons were found in genetic models with band heterotopia (15, 21). Last, ectopic neurons displayed various degrees of simplification of dendritic arbors in all models (17, 18, 20), as well as an apparent reduction of spine density (19) in line with our results. Therefore, if axonal development is relatively spared from the consequences of altered laminar placement, our observations confirm that the eventual positioning of neocortical neurons appears instrumental to the subsequent morphogenesis of dendrites, and synapses.

The mechanism responsible for impaired dendritic maturation of ectopic neurons remains to be clarified. Cell-extrinsic mechanisms have been shown to play major roles in dendrite morphogenesis (22) which can further influence cell-intrinsic mechanisms. Extrinsic cues include numerous secreted factors (23, 24) and contact-mediated cues (25, 26), as well as neuronal activity (27). Ectopic neurons are clearly not exposed to the same microenvironment as normally positioned layer V neurons and they may not receive the same inputs as if positioned in layer V, which may influence negatively dendrite development. Recent studies have shown that the development of the dendritic arbors and assembly of excitatory synpases can occur in the absence of glutamatergic neurotransmission (28 – 33) even though one of these studies reported that both the dendritic arborization as well as the linear spine densities were reduced in its absence (32). A previous study on the Dcx-KD model showed that most ectopic neurons displayed a delayed maturation of GABA-mediated signaling (34). Since early developing GABAergic inputs from cortical interneurons have been shown to control both inhibitory and excitatory synaptogenesis (35), defective synaptogenesis in ectopic neurons might result from a defective GABAergic signaling in these neurons.

Our experiments also highlighted a novel role for Dcx in regulating the dendritic and synaptic development of cortical neurons independently of their laminar placement. Both in ectopic and properly placed neurons, knockdown of Dcx lead to a decreased number of dendritic branches when compared to Dcx^+^ neurons in the same position. These results are in line with studies carried out in cultured neurons where Dcx knockdown or mutated forms of human Dcx overexpression resulted in impaired development of the dendritic arbor (36 – 38). Because dendritogenesis relies heavily on microtubules stabilization, Dcx effect could be mediated through its function as a microtubule associated protein (39 – 42). Indeed, Dcx stabilizes microtubules by linking adjacent protofilaments and counteracting their outward bending in depolymerizing microtubules (43, 44). Moreover, its localization at the distal part of growing neuronal processes places Dcx at a key spot to regulate dendritic branching (45 – 47). Formation of a new dendritic branch can be achieved either through sliding of pre-assembled microtubules from the soma into the dendrite or through microtubule nucleation, a process leading to *de novo* polymerization of microtubules (48). Since Dcx has been shown to nucleate microtubules *in vitro* and *in vivo*, microtubule nucleation could be the process through which Dcx promotes dendritic ramification of cortical neurons (39, 43, 44).

In addition to Dcx involvement in dendritogenesis, we also found a significant enlargement of spine head and neck diameters in Dcx-pKD neurons, suggesting that Dcx acts as a regulator of dendritic spine shape. This finding is in agreement with a previous report identifying Dcx as the downstream effector of Npas4 and Mdm2 which regulate spine formation in newborn neurons of the olfactory bulb (49). Spine formation and plasticity rely heavily on the actin cytoskeleton to control spine shape. In developing neurons, Dcx regulates filamentous actin by interacting with neurabin II (spinophilin) thus cross-linking the actin and microtubules cytoskeleton (50 – 52). Furthermore, although spines have been considered devoid of microtubules for a long time, several studies have now underlined the presence of dynamic microtubules in these structures (53, 54). Microtubules polymerize transiently into spines and regulate their morphology as well as synaptic plasticity through a cytoskeletal cross-talk with actin (55 – 57). Because microtubule invasions seem to specifically target spines that are undergoing activity-dependent changes, one hypothesis is that this process helps to regulate synaptic plasticity through molecular cargo transport to the synapse (58, 59). Overall, Dcx many interactions with the cytoskeleton are likely to contribute to its regulation of dendritic development, spine morphology and synaptogenesis.

Lastly, our paper identifies laminar misplacement resulting from migration failure as a critical factor holding up the development of cortical neurons. In humans, migration failure during corticogenesis leads to malformations of cortical development that are often associated with epilepsy and intellectual disability (2, 60). These symptoms are primarily thought to arise from the structural alterations created by malformations as well as aberrant network connections. However, as these malformations are formed by groups of ectopic neurons, the dendritic and synaptic deficits described above could participate to the etiology of the clinical symptoms. In fact, dendritic shape impairment during development has been linked to lower cognitive capacities later in life (61, 62). Moreover, spine alterations and synaptic dysfunctions are well-known mechanisms underlying brain deficits that are known collectively as synaptopathies (63). Finally, since Dcx also seems to regulate some aspects of neuronal development, this study may be relevant for a migration disorder arising from Dcx mutations, called subcortical band heterotopia (SBH) (64 – 68). Patients with SBH suffer from drug-resistant epilepsy and intellectual disability. According to our results, laminar misplacement and Dcx deficiency could contribute to the pathophysiology of SBH by acting synergistically to alter the maturation of ectopic neurons.

## METHODS

### Animals

Animal experiments were performed in agreement with European directive 2010/63/UE and received approval from the French Ministry for Research, after ethical evaluation by the institutional animal care and use committee of Aix-Marseille University [protocol number: 2015040809544569_v2 (APAFIS#436)].

All experimental and control animals were generated by in utero electroporation at embryonic day 15 (E15) as described previously (69). Briefly, timed pregnant Wistar rats (Janvier) were anesthetized either with a mix of ketamine (Imalgene 1000 at 100 mg/kg) and xylazine (Rompun 2% at 10 mg/kg) or with sevoflurane 4%. Sevoflurane anesthetized rats received buprenorphine (Buprecare at 0.03 mg/kg) 30 minutes before surgery. Uterine horns were exposed and a mix of plasmids (see below) and fast green was injected by pressure (PV 820 Pneumatic PicoPump; World Precision Instruments, Sarasota, FL) in the lateral ventricles of embryos through pulled glass capillaries (Drummond Scientific). Electroporations were accomplished by discharging a capacitor with a sequencing power supply (BTX ECM 830 electroporator; BTX Harvard Apparatus, Holliston, MA). The voltage pulse (40V) was discharged across tweezer-type electrodes (Nepa Gene Co, Chiba, Japan) pinching the head of each embryo through the uterus.

To induce ectopic positioning of neurons, we electroporated plasmids encoding shRNAs targeting the 3’ untranslated region (3’UTR) of *Dcx* (mU6pro-3’UTRhp; gift from Joe Loturco (9)). For control, we used plasmids encoding ineffective shRNAs with 3 point mutations to create mismatches (mU6pro-3’UTR-mismatch; gift from Joe Loturco (9)). Whereas the former construct efficiently knocks down Dcx expression in vivo, the latter construct is ineffective in causing Dcx knockdown (9). The coelectroporation of plasmids encoding the mCherry (pUbiquitin-mCherry; gift from I. Medyna) or the enhanced green fluorescent protein eGFP (pCAG-GFP; Addgene #11150) allowed visualization of electroporated neurons. Post-synaptic sites were genetically labelled with a conditional Floxed-Stop plasmid expressing PSD-95-GFP (pCALSL-PSD-95-GFP; gift from Hiroshi Kawazaki (70)). Recombination of this plasmid was induced by the co-electroporation of a 4-hydroxy-tamoxifen (4-OHT)-activable form of Cre recombinase (pCAG-ERT2CreERT2; Addgene #13777). To generate Dcx-pKD animals, the sequences targeting the *Dcx* 3’UTR from the mU6pro-3’UTRhp were cloned into the conditional Floxed-Stop miR-30 plasmid (pCALSL-miR30; Addgene # 13786) giving rise to the pCALSL-mir30-3’UTRhp. Mismatch sequences identical to those of the control plasmid mU6pro-3’UTR-mismatch were cloned in the pCALSL-miR30 to generate the control plasmid pCALSL-mir30-3’UTR-mismatch. All plasmids were injected at a concentration of 1 μg/μl excepted the pUbiquitin-mCherry and pCAG-GFP which were injected at a concentration of 0.5 μg/μl. Cre-ERT^2^-dependent recombination was carried out by injecting pups with a single intraperitoneal shot of 4-OHT (Sigma, 2 mg/mL diluted in corn oil (Sigma), 120-160 μg / pup) at P1.

### Electrophysiological recordings

Animals were anesthetized and then decapitated. The brain was removed rapidly and transverse 300 μm thick slices were cut using an LeicaVT1200S tissue slicer in a solution containing the following (in mM): 110 choline, 2.5 KCl, 1.25 NaH2P04, 25 NaHCO3, 7 MgCl2, 0.5 CaCl2, and 7 D-glucose, 300±10 mOsm, pH 7.4 (4°C). Slices were transferred for rest (at least 1 hr) at room temperature in oxygenated normal artificial cerebrospinal fluid (ACSF) containing the following (in mM) : 126 NaCl, 3.5 KCl, 1.2 NaH2PO4, 26 NaHCO3, 1.3 MgCl2, 2.0 CaCl2, and 10 D-glucose, 300±10 mOsm, pH 7.4.

Individual slices were then transferred to the recording chamber, where they were superfused with ACSF at 30±1°C at a rate of 2-3ml/min. Electropororated /non electroporated cells in layer 5 were recorded in whole cell configuration in voltage-clamp mode with borosilicate glass micropipette (Resistance 6-8mΩ) containing (in mM): 130 CsGlu, 10 CsCl, 0.1 CaCl2, 1.1 EGTA, 10 HEPES, 4 Mg2+ATP, 0.3 Na+GTP, pH 7.22, 279 mOsm. For miniatures recordings, 1μM TTX (Tocris, UK) was added in ACSF. Biocytin (0.5%, Sigma, USA) was added to the pipette solution for post hoc reconstruction.

Whole-cell measurements were made using a Multiclamp 700B amplifier (Axon Instruments, Molecular Devices, USA) with a Digidata 1440A (Molecular Devices) and acquired using pCLAMP 10.3 software (Molecular Devices, USA). Signals were analyzed off line using MiniAnalysis 6.0.7 (Synaptosoft, USA) by the experimenters blind to experimental conditions.

### Immunostaining

Animals were transcardially perfused with AntigenFix (Diapath) at P14-15. Their brains were then removed and post-fixed in AntigenFix for 24 hours before they were sectioned (100 μm) with a vibratome (Leica VT 1000S) and processed for immunohistochemistry as free-floating sections. The mCherry signal was amplified using an anti-RFP rabbit antibody (1/1000, Rockland). Sections used for electrophysiology were fixed overnight in AntigenFix before biocytin was revealed with Alexa647-conjugated-streptavidin (1/200, Jackson ImmunoResearch). In all cases, sections were incubated with antibodies for 72 hours at room temperature to increase antibody penetration into the tissue. Sections were counterstained with Hoechst (Thermo Fisher, 1/1000) and mounted in Fluoromount (Thermo Fisher).

### Fluorescence microscopy

All images were acquired as 3D stacks using a 0.3 μm z-step on a Leica TCS SP5 X confocal microscope, unless stated otherwise. For the morphological analyses of ectopic / layer V and Mismatch / Dcx-pKD dendritic trees, neurons were imaged using a 63x oil objective, roughly placing the soma at the center of the image. During the same imaging session, dendrites were imaged by using the same objective and adding a 4.5 numerical zoom. Only dendrites within 5-80 μm of the soma were imaged. When two channels were used, acquisitions were always sequential to avoid cross-talk. After electrophysiological recordings, a 40x oil objective was used to image the biocytin-filled neurons.

Images used for reconstructions of ectopic Dcx^+^ and Dcx-KD neurons were acquired using a Zeiss LSM 510 confocal microscope with a 20x objective and 0.5 μm z-step.

### Electron microscopy

PSD95-GFP electroporated P15 rats were deeply anesthetized and transcardially perfused with AntigenFix (Diapath) /0.3% glutaraldehyde (Sigma). Brains were dissected out and post-fixed overnight in Antigen Fix, at 4°C. 120 μm coronal sections were prepared using a vibratome. PSD95-eGFP electroporated sections were selected under an epifluorescence microscope, cryoprotected in 25% sucrose in PBS overnight at 4°C and submitted to two cycles of freezing on dry ice and thawing, for permeabilization. Selected sections were incubated in blocking buffer (PBS, 5% BSA, 0.1% cold water fish skin (CWFS) gelatin, 10% normal goat serum (NGS), 15 mM NaN3 pH7.4) for 1 hour at room temperature, incubated with a chicken anti-eGFP antibody (1/1000, Aves) in antibody buffer (PBS, 0.8% BSA, 0.1% CWFS gelatin, 5% NGS, 15 mM NaN3 pH7.4) for 72 hours, at 4°C, washed and further incubated with a gold-coated IgG goat anti-chicken antibody (1/50, Aurion, “Ultra small 0.8 nm”, Cat N° 800.244) in the same buffer for 16 hours at 4°C. Sections were washed and post-fixed in PBS/2.5% glutaraldehyde at room temperature for 30 min. The sections were treated with the Aurion R-Gent silver enhancement kit (Electron Microscopy Sciences) and post-fixed with 0.5 % osmium tetroxide in PBS at room temperature for one hour. The sections were dehydrated in a graded series of ethanol from 30% to 70% and stained overnight in 2% uranyl acetate in 70% ethanol at 4°C. The sections were further dehydrated in a graded series of ethanol from 70 % to 100%, washed with propylene oxide, embedded in Araldite resin. 80 nm sections were made with an ultramicrotome (Leica, EMUC7), analyzed with an electron microscope (Zeiss EM 912). Images were acquired with a digital camera (Bioscan 792).

### Image analysis

All image analyses were performed with the scientist blinded to the experimental conditions.

Radial orientation of neurons was defined as the apical dendrite making an angle of 90° ± 20° with the cortical surface.

Neurons were reconstructed tree-dimensionally using Neurolucida software version 10 (MBF Bioscience). The digital reconstructions were analyzed with the software L-Measure to measure the number of primary branches and the total number of ramifications of each neuron (71).

PSD-95-GFP^+^ puncta were manually counted on the 3D stack using the Cell counter plugin of FIJI. The length of the corresponding dendrite was also measured in FIJI. The linear density of PSD-95-GFP^+^ puncta was calculated as the ratio of these two measurements.

Dendritic spine density and morphology were analyzed from 2D images generated by maximum intensity projection of the 3D stacks in FIJI and converted to RGB images. The length, head width and neck width of dendritic spines were then manually traced on SynPAnal (72). Only spines clearly protruding from the dendritic shaft were reconstructed. A 700 nm wide circle was then generated around each spine head and PSD-95-GFP^+^ puncta were detected using a fluorescence threshold to count the proportion of spines containing puncta. This detection was manually edited to remove false positive.

### Synaptosome preparation and Western blot analysis

Cortex from P10, 15 and P20 rat were homogenized in Syn-PER^TM^ synaptic protein extraction reagent (ThermoScientific) complemented with protease and phosphatase inhibitors (Pierce, ThermoScientific) in a Dounce homogenizer (1ml/100 mg of tissue) at 4°C. The homogenates were centrifuged at 1000 g for 10 min at 4°C and the supernatants further centrifuged at 15000g for 20 min at 4°C. The supernatants (cytosolic fractions) were collected and the pellets (synaptosome fractions) were resuspended in 200 μl of Syn-PER reagent, with protease and phosphatase inhibitors (Pierce, ThermoScientific). Proteins from cytosolic fractions and synaptosomes samples were quantified using a bicinchronic acid (BCA) Protein Assay Kit (Pierce, ThermoScientific). 20 μg of proteins were loaded par well, subjected to 7.5 % sodium-dodecyl sulfate-polyacrylamide gel electrophoresis (SDS-PAGE) (Criterion™ TGX^TM^ precast gels, Bio-Rad Laboratories) and transferred to Immun-blot LF PVDF membranes (Bio-Rad Laboratories). Membranes were blocked for 1 hour and incubated overnight with rabbit polyclonal anti-Dcx (1/1 000, Abcam, Cat No AB18723) and rabbit monoclonal anti-PSD95 (D74D3) (1/1000, Cell Signaling, Cat No 3409S) antibodies. Membranes were then incubated with an Alexa Fluor 647 goat anti-rabbit antibody (1/500, Invitrogen) for 1 hour. Image acquisitions were performed with a CDD camera (GBox, Syngene). The membranes were incubated in stripping buffer (62.5 mM Tris-HCl pH6.8, 2% Sodium Dodecyl Sulfate (SDS), 100 mM β-mercaptoethanol) at 55°C for 30 min, blocked and re-incubated with a mouse anti-α-tubulin (DM1A) antibody (1/5000, ThermoScientific, Cat No 62204) for 4 hours. Membranes were washed and incubated with an Alexa Fluor 488 goat anti-mouse antibody (1/500, Invitrogen, Cat No A11001) for 1 hour and imaged.

### Statistics

All statistical analyses were performed using Prism 6 (Graphpad). Normality of the data distributions was systematically tested using d’Agostino & Pearson test and Shapiro-Wilk test. Comparison of groups was subsequently tested with unpaired t-tests for normal data sets or Mann-Whitney tests for non-normal data sets. Two-sample Kolmogorov-Smirnov tests were used to compare distributions of data obtained from electrophysiological experiments. All values are given as mean ± SEM. All tests were two-tailed and the level of significance was set at P<0.05. Statistical power was checked in all experiments to ensure sample sizes were adequate. n refers to the number of cells, except for spine morphology analyses where it refers to the number of spines.

### Data availability

The data that support the findings of this study are available from the corresponding authors upon request.

## ACKNOWLEDGEMENTS

We thank the animal facility (PPGI, INMED, Marseille), the imaging facility (INMAGIC, INMED, Marseille) and the electron microscope facililty (IBDM, Marseille), Dr Roman Tyzio for his help in the preliminary electrophysiology experiments, Laetitia Weinhard for her help in the preliminary histology experiments, Benoît Boulan for RNAi plasmid clonings, Jean-Christophe Vermoyal for Neurolucida neuronal reconstructions, and Eric Danielson for his help with spine analyses. We thank Fiona Francis and Sonia Garel for critical discussions and comments on the manuscript. This work was supported by La Fondation pour la Recherche Médicale (#FDT20160435216, F.M.), the French National Agency for Research (SAMENTA, #19012012, A.R.) and the European Community 7th Framework programs (DESIRE, Health-F2-602531-2013, AR; DECIPHER, #ANR-15-NEUR-0001-03, A.R.)

## AUTHOR CONTRIBUTIONS

F.M., F.W., J.B.M. and A.R. designed and conceived the experiments. E.B. and F.S. performed in utero electroporations. L.F. and F.W. performed the cloning and preparation of plasmids. H.K provided the PSD-95-GFP plasmid. F.W. prepared the synaptosomes and performed the Western blot analyses. F.M. and F.W. performed morphological analyses. G.C. and F.W. performed electron microscopy experiments. V.P. and S.S. performed electrophysiological experiments. F.M, S.S, V.P., F.W., A.R. and J.B.M. analyzed the data. F.M., F.W., A.R., and J.B.M. wrote the manuscript.

## COMPETING FINANCIAL INTERESTS

The authors declare no competing financial interests.

## MATERIALS & CORRESPONDENCE

Correspondence and requests for materials should be addressed to F.W. and J.B.M.

